# Cortical elasticity determines the geometry of prospective mesoderm constriction in *Drosophila melanogaster*

**DOI:** 10.1101/102277

**Authors:** Konstantin Doubrovinski

## Abstract

An animal embryo begins its life as a ball of epithelial cells. In the course of development, invariably, this cellular ball will undergo a process of gastrulation to form a multilayered structure with the different germ layers designated to form organs with different shapes and functions. In the fruit fly *Drosophia melanogaster*, gastrulation begins with the constriction of mesodermal cells that make up a rectangular domain in the ventral part of the embryo. A remarkable aspect of this morphogenetic event is its anisotropy - the mesoderm constricts much more along one axis than along the other. In this paper we propose an explanation of this observed anisotropy. Specifically, we show that tissue contraction must be anisotropic, provided that the tissue is elastic and that the contractile domain is elongated (e.g. rectangular as opposed to square). This conclusion is generic in the sense that it does not depend on the specific values of model parameters. Since our recent study demonstrated that embryonic tissue is elastic on a developmentally relevant time-scale, it appears likely that the anisotropy of mesoderm contraction is an elastic effect. Our model makes a number of specific predictions that appear in close agreement with the available data.

## 2 Introduction

In the fruit fly *Drosophila melanogaster*, development starts with thirteen nuclear divisions. In the course of these nuclear cleavages the embryo remains a syncytium, with all nuclei sharing common cytoplasm within a large, half-millimeter long cell. By the tenth mitotic nuclear division most nuclei have migrated to the surface of the embryo [1]. After the thirteenth mitotic cycle, in a process called cellularization, membranes protrude from the surface of the embryo to engulf the nuclei and thereby partition the embryo into some six thousand distinct cells (blastocytes) [1, 2]. In the course of cellularization, the newly-forming cells acquire epithelial character, with the various polarity markers, organelles, and sub-cellular structures arranged as is characteristic of epithelia of many organs[3]. Apical domains of the blastocytes face outwards, forming the surface of the embryo. Cellularization is followed by the process of gastrulation, whereby a single epithelial sheet gives rise to a multilayered structure [4, 5]. Gastrulation initiates when a set of cells located in a rectangular domain in the ventral part of the embryo constrict apically. These contractile cells have mesodermal character that is conferred by the expression of transcription factors *snail* and *twist* [6]. As mesodermal cells constrict apically, they become wedge-shaped and elongate along the baso-lateral axis. Apical constriction of the mesodermal cells is widely believed to be driven by myosin-generated stresses in the apical domains of those cells. In accordance with this, myosin concentration increases dramatically in the constricting cells concomitantly with the onset of apical constriction. Following apical constriction, the surface of the embryo forms a furrow at the ventral midline. The furrow deepens and closes off; in this way, mesoderm is brought into the interior of the embryo.

The focus of the present study is the initial phase of gastrulation when mesodermal cells constrict apically, before the surface begins to fold. Notably, as mesodermal cells shrink, the rectangular mesodermal domain contracts strongly along its shorter axis and much less so along the longer axis. For brevity, we will refer to the length of the mesodermal domain along the mediolateral axis as the “width”, and we will refer to the length of the mesodermal domain along the anteroposterior axis as the “length”. Thus, the length to width ratio increases drastically in the course of the constriction of the mesoderm, see schematic in Figure 1a. It is well established that mesoderm constriction in *Drosophila melanogaster* is not accompanied by cell rearrangements. Thus, the length to width ratio of individual mesodermal cells must increase in the course of tissue contraction, as is indeed seen to be the case, see e.g. [7].

In the present work we propose a simple mechanistic explanation of the asymmetry of constriction of the mesoderm. The proposed mechanism principally relies on our recent measurements of material properties of the embryonic tissue in *Drosophila melanogaster*. We shall see that the proposed mechanism is generic (does not require specific assumptions about model parameters), and, at the same time, predictive, since it relates a number of independent and seemingly unrelated experimental observations about the dynamics of mesoderm constriction.

**Figure 1.**
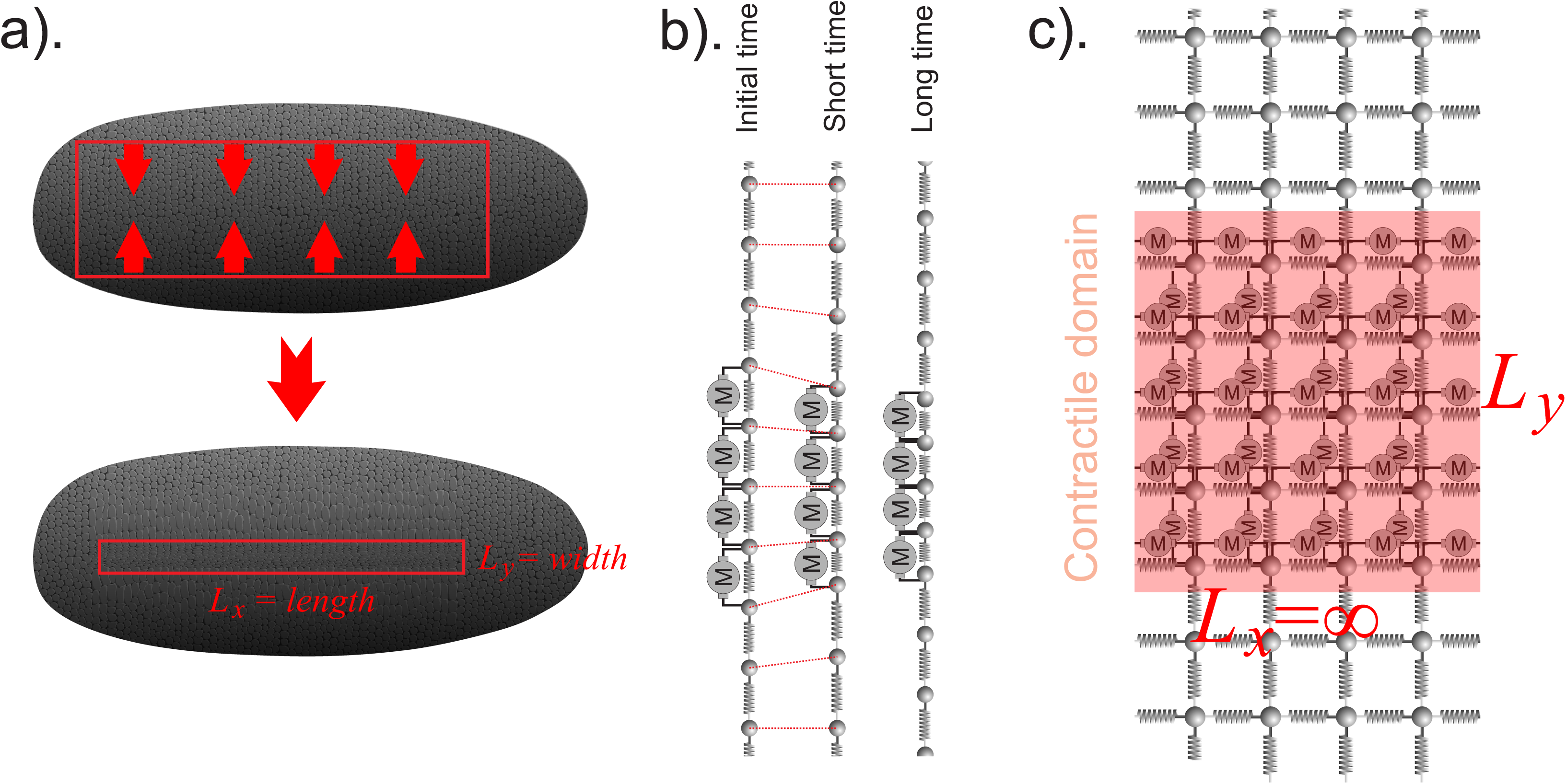
Schematic illustrating the major ingredients of the model. a). Schematic of a gastrulating embryo. Anteroposterior axis is horizontal, medialateral axis is vertical. Approximately ten minutes after the onset of gastrulation, mesoderm constricts (bottom panel). Mesoderm contraction is much stronger along the medialateral (vertical, “width”) axis than along the perpendicular antoroposterior (horizontal, “length”) direction. b). Schematic of the onedimensional limit of the model. Elastic elements are illustrated as springs, active forces are represented by “motors” (round circles with letter “M”) whose two force-excerting “arms” attach to adjacent “beads”. Beads are assumed to feel viscous drag from the ambient environment when moving. Note that it is assued that forces axcerted by motors are constant in time and space. c). Schematic of the two-dimensional model. All symbols are as in b). The schematic illustrates the case when contractile domain is infinite along its length. Then, by symmetry, deformation does not vary along the horizontal direction.

## 3 Results

The aim of the present paper is to present a model that explains the geometry of mesoderm constriction in *Drosophila melanogaster*. Instead of starting with a formal statement of the model and formal treatment of the corresponding equations, we begin by stating the major conclusions and give an intuitive argument leading up to those conclusions. Thereafter, a more formal treatment is given.

We model the surface of the embryo as a flat (two-dimensional) elastic sheet. For the sake of simplicity we may assume that the sheet is infinite (occupying the whole *x-y* plane) and thus disregard any boundary effects. Our model sheet will contain a rectangular contractile domain assumed to have horizontal and vertical dimensions *L_x_* (length) and *L_y_* (width) respectively, with *L_x_* > *L_y_* (see schematic Figure 1a). Note that we are here assuming that both dimensions of the contractile domain are finite. In order for this domain to contract it must carry contractile stress (or “tension”). Loosely speaking, this domain “wants to shrink”. We assume the stress within the contractile domain is uniform (having the same magnitude everywhere) and isotropic (same in any direction). Outside of the contractile domain the stress is absent. Note that in the real embryo, contractile stress is generated by myosin motors. Finally, we assume that our elastic sheet feels viscous drag from its ambient environment. This viscous drag may, for example, result from the interaction of cell surfaces with the cytoplasm. Suppose now that stress in the contractile domain is absent before time zero, and is subsequently turned on to its full extent. We claim that the contractile domain will start to shrink predominantly along its shorter dimension. Thus, the length to width ratio of the contractile domain will start to increase upon the appearance of contractile stresses (corresponding to the onset of gastrulation). Moreover, as we show below, the aspect ratio will not return to its initial value at long time, and, hence, the domain will remain anisotropic as long as contractile stresses persist.

To intuitively explain the first claim, we will consider a number of physical models in order of increasing complexity. In this way, our claim about the asymmetry of contraction under the above assumptions will be made more intuitive. First, let us consider the situation depicted in the schematic in Figure 1b. We are considering an infinite chain of springs connected in series. These springs are assumed to be linearly elastic and to have the same spring constant. Any two adjacent springs meet at a node that is represented by a spherical bead. If one of these beads moves, it is assumed to experience drag from its ambient environment which is proportional to the speed of the bead. Finally, the springs located within a certain region are assumed to be under tension. More specifically, one can picture “motors” that apply forces to adjacent beads causing those beads to (effectively) attract (see schematic in Figure 1b). In what follows, we will refer to the forces applied by the motors as “active” forces. We assume that these active forces stay constant after *t* = 0. Taken together, at any point in time, the force on any given bead has the following three contributions: the forces from the two adjacent springs, the forces from the two “adjacent motors”, and viscous drag from the ambient fluid. Note that the forces from the two adjacent springs depend exclusively on the relative positions of the given bead with respect to its two nearest neighbors (since spring force is determined exclusively by the extension of a spring). Suppose now that motors are off initially, and turn on instantaneously at time zero. Clearly, the contractile domain will gradually shrink. At short times the deformation will be confined to the vicinity of the boundaries of the contractile domain, Figure 1b. Subsequently, beads further and further away from the contractile domain boundaries will start to move. This situation may be compared to diffusion. If a diffusible substance is continually supplied to a small region of space, the substance will spread gradually, reaching further and further away from the region where it is being deposited. This analogy may be made precise since, in fact, mathematically, our one-dimensional elasticity problem is precisely equivalent to a one-dimensional diffusion problem. (In this way, active motor-driven force in our mechanical problem is analogous to the source of the diffusible substance, while the displacement of a bead at a given position in space is analogous to the concentration of the diffusible substance at the respective spatial position). Finally, let us make one last observation about this one-dimensional toy model. Consider any given bead in the *interior* of the contractile domain. The force from the left adjacent motor is always the same and opposite as the force from the right adjacent motor. In this way, *active* forces on any interior node cancel and do not influence the dynamics. Hence, nothing would change if we removed all motors and instead replaced them with two “point forces” applied to the left-most and the right-most beads of the contractile domain and directed towards the right and left respectively. Note, however, that the total force, (i.e. the sum of all three above-mentioned force contributions) is, in general, not zero at an interior node.

Now, let us consider a two-dimensional analogue of the one-dimensional toy model from the previous paragraph. Consider a network of springs arranged in a regular grid of squares, as shown in the schematic Figure 1c. For simplicity of the argument we assume this grid to be infinite in both dimensions. Furthermore, the vertices of the grid correspond to points of the coordinate plane with integer coordinates. Suppose now that the contractile domain has infinite length and finite width, i.e. the contractile domain is the region −∞ *< x <*∞, −*L_y_* /2*< y < L_y_*/2, see Figure 1c. A moment of thought shows that the dynamics in this quasi-two-dimensional model are the same as in the one-dimensional case considered in the previous paragraph. This is because, by symmetry, all nodes will displace exclusively along the vertical direction. Thus, the deformation is one-dimensional, being the same along any one vertical section of the plane.

Finally, consider the case when the contractile domain is finite in both directions, i.e. occupies the region −*L_x_/*2 *< x < L_x_/2*, −*L_y_*/2 < y < *L_y_*/2. Suppose *L_x_* >> *L_y_*, i.e. the length is much larger than the width. Let us first focus our attention on the portion of the contractile domain in the vicinity of its center, i.e. on some region −ϵ < x < ϵ, −L_y_/2 < y < L_y_/2, with e much smaller than Lx. Now recall from the above discussion that the deformation propagates away from the boundary of the contractile domain in a diffusive manner (i.e. on a finite time-scale). Thus, in the considered portion of the contractile domain, the dynamics will not be influenced by the presence of the left-most or right-most boundaries of the domain until the deformation has had time to propagate from those boundaries all the way to the center. Hence, in the central portion of the domain, the contraction will initially happen predominantly along the shorter dimension.The discussion in the previous paragraph raises the following question: will the aspect ratio of the domain return to its initial value if given enough time? Below we present a formal treatment showing that the answer is “no”: the domain will remain constricted anisotropically, and this is true for almost all values of the model paameters. It is harder to give an intuitive account for this behavior than to intuitively explain the initial transient dynamics, but we will show this formally later in the text.

Taken together, we have described a model that we will show can explain the asymmetry of mesoderm constriction. First, however, we will consider a number of model predictions. Firstly, if the contractile domain were symmetric (e.g. round or square) it would constrict symmetrically. Such an experiment has in fact been done in a classical paper by Maria Leptin and Siegfried Roth [8]. There, patches of mesodermal tissue (either nuclei or cytoplasm) were transplanted into mutant embryos that lacked mesoderm. It was shown that the geometry of contraction of mesodermal patches was independent of the position and orientation of those patches within the embryo and was only dependent on the geometry of the patch. In particular, roundish patches remained round, whereas elongated patches contracted predominantly along their shorter dimension. These observations are in agreement with and serve as evidence for our interpretation. Additionally, our model correctly predicts the outcome of laser ablation experiments. Specifically, it was previously shown that when the surface of the mesoderm is laser ablated to create an initially round hole, the hole will expand predominantly along the longer axis of the embryo [9]. This “anisotropy” of the response to ablation appears to increase as gastrulation proceeds: immediately after the onset of gastrulation the response to ablation is approximately “isotropic”, becoming gradually more “anisotropic” with time, see [9]. These observations are in full accordance with our model. To see this intuitively, let us again consider the idealized model in Figure 1c with a contractile domain that extends infinitely in one direction. After the onset of contraction, the domain will shrink along the shorter vertical direction and not at all along the perpendicular horizontal direction. Now, consider what would happen if one were to remove a single vertical spring-edge in the interior of the contractile domain after contraction sets in, see Figure 3b’. This vertical spring-edge has two adjacent vertical edges. If the contractile domain had already undergone some contraction, those adjacent vertical neighbors would have shrunk to become shorter than their preferred rest length. Spring forces in those adjacent spring edges would tend to expand those adjacent springs, thus counteracting the expansion of the hole along the vertical direction. Now, instead, consider what would happen if a horizontal spring-edge is removed. The adjacent horizontal spring-edges are under no compression. Thus, elastic forces in those springs will counteract the expansion of the “hole” to a lesser extent than was the case with the vertical spring-edge. In this argument, we have been assuming that *active* forces on every spring in the interior of the contractile domain are the same on every edge of the network. This is to say that our schematic “motors” that serve to shrink the contractile domain are all the same and do not change with time, as has been assumed throughout. Note that in order to correctly interpret the outcome of the ablation experiments it is absolutely necessary to distinguish between “total stress”, “elastic stress” and “active stress”. Elastic stress is due to the springs. Active stress is due to the motors. Total stress is the sum of the two. Ablation experiments measure total stress which becomes increasingly anisotropic in the course of the contraction. Active stress, however, remains isotropic throughout the course of the dynamics. Previously, the anisotropy in response to ablation was interpreted as evidence for anisotropy of the active stress [9]. Our model provides a significantly simpler interpretation: total stress must become anisotropic if the domain is asymmetric (and if the tissue is elastic), whereas active stress need not be anisotropic. Also, in our proposed model, the asymmetry of contraction is the cause, not a consequence, of anisotropic total stress as measured by laser ablation. Taken together, we have seen that two additional key observations about the dynamics of mesoderm contraction are readily explained by our proposed minimal model.

We shall next turn to a more formal mathematical treatment of the model sketched above. We considered the surface of the embryo as a flat (twodimensional) elastic sheet with several properties. (1) The sheet is immersed in a viscous environment; thus, as the sheet deforms, it will experience viscous drag from its ambient environment. (2) The region of the sheet that corresponds to the mesoderm is subjected to contractile stress. (3) The stress is uniform and isotropic in the mesoderm and zero in the rest of the tissue. Combinind these assumptions with standard equations of linear elasticity (see [11], in particular Equations 13.4 therein) we have:

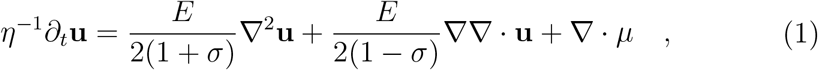

where **u** is the in-plane deformation of the sheet, *η* is the coefficient describing viscous drag between the sheet and its viscous environment, *E* is Young’s modulus (more specifically, this quantity is *Eh*, where *E* is the three-dimensional Young’s modulus, and *h* is the thickness of the sheet), *σ* is the Poisson’s ratio, and *μ* is the stress within the contractile mesodermal domain. We set *μ* = *ϕ***I**, with **I** being identity matrix, and *ϕ*, the magnitude of stress, being constant in the mesodermal domain and zero outside. With a slight abuse of notation we denote the constant magnitude of stress within the mesoderm by *ϕ* as well. Our model is schematically illustrated in Figure 1c. The elastic sheet is represented by a spring network with nodes illustrated as spherical beads. Those beads may be thought of as having a finite hydrodynamic radius, implying that those beads experience viscous drag in the course of deformation. Note that in the interest of simplicity the illustration was made somewhat misleading: a network squares is not isotropic (the stiffness is not the same in all directions), whereas Equation (1) describes an isotropic elastic material. To re-iterate, our description is exceedingly simple: an elastic sheet feels viscous drag from its ambient environment (cytoplasm) and (myosin-generated) contractile stress is constant and isotropic within the rectangular (mesodermal) domain and zero elsewhere.

We begin the analysis of our minimal model given by Equation (1) by non-dimensionalizing the equation. Dividing throughout by *ϕ*, rescaling length by the width of the contractile domain *L_y_*, and rescaling time by *ηL_y_*/*ϕ*, we obtain the re-scaled equation in the same form as Equation (1), except with *η* and *ϕ* set to one, and Young’s modulus E replaced by the dimensionless ratio *E*/*ϕ*. Taken together, our equation has the following three dimensionless parameters: the ratio of Young’s modulus to stress magnitude *E*/ϕ, Poisson’s ratio *σ*, and the length to width ratio of the contractile domain *L_x_/L_y_*.

To analyze the dynamics of our minimal model (1) we performed numerical simulations, see Figure 2. We simulated an isotropic, flat, elastic sheet whose dynamics are overdamped and driven by contractile stresses in a rectangular domain. The size of the simulated sheet is taken to be much larger than the size of the contractile domain in order to avoid finite-size effects. Implementation details are given in Appendix A. Our simulations indicate that if the contractile domain is chosen to be asymmetric (i.e. length larger than width), constriction is found to be stronger along the shorter axis of the domain, see Figure 2. This effect is not transient - anisotropy persists at long times.

**Figure 2.**
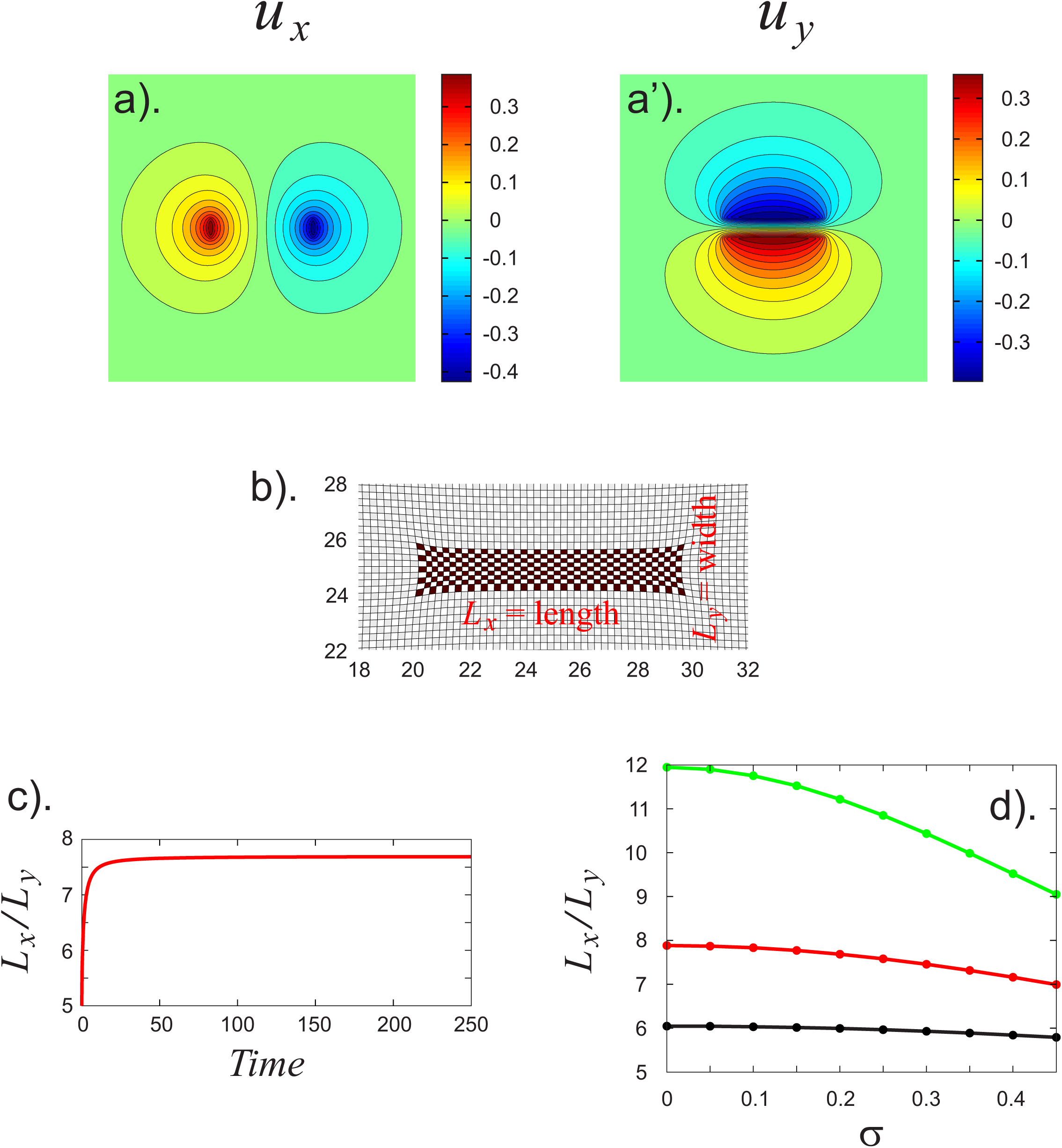
Simulations showing the deformation of a flat elastic shell by unifrom isotropic stresses confined to a rectangular contractile domain. a)-a’). Color plots of the deformation of an elastic shell in mechanical equilibrium. a) and a’) show *x*- and *y*-components of the deformation respectively. Parameters are: *E* =1, *η* =1, *σ* = 0.2, *ϕ* = 0.5, *L_x_* = 10, *L_y_* = 2, (see main text for notation), domain size is 50x50. For readability only the noticably deformed middle portion of the domain with a side length of 30 is shown. b). Same as a), except the deformation is illustrated as distortion of a uniform grid of squares. Note that squares in the middle of the domain are distorted into rectagles with their long axis along the long axis of the contractile domain. c). Length to width ratio (*L_x_*/*L_y_*) of the contractile domain as a function of time for the simulation that produces equilibrium state shown in a) and b). Note that asymptotically, the ratio is larger than it is initially. If this curve were normalized by the value at *t* = 0, the resulting function would show apprximate aspect ratio of the “square” in the center of the contractile domain in b). c). Assymptotic (equilibrium) aspect ratio of the contractile domain as a function of Poisson’s ratio (*x*-axis) and contractile stress. All parameters (exceps the ones that are varied) are as in a). Black, red, and green curves correspond to *ϕ* = 0.5, *ϕ* = 0.75, and *ϕ* =1 respectively.

**Figure 3.**
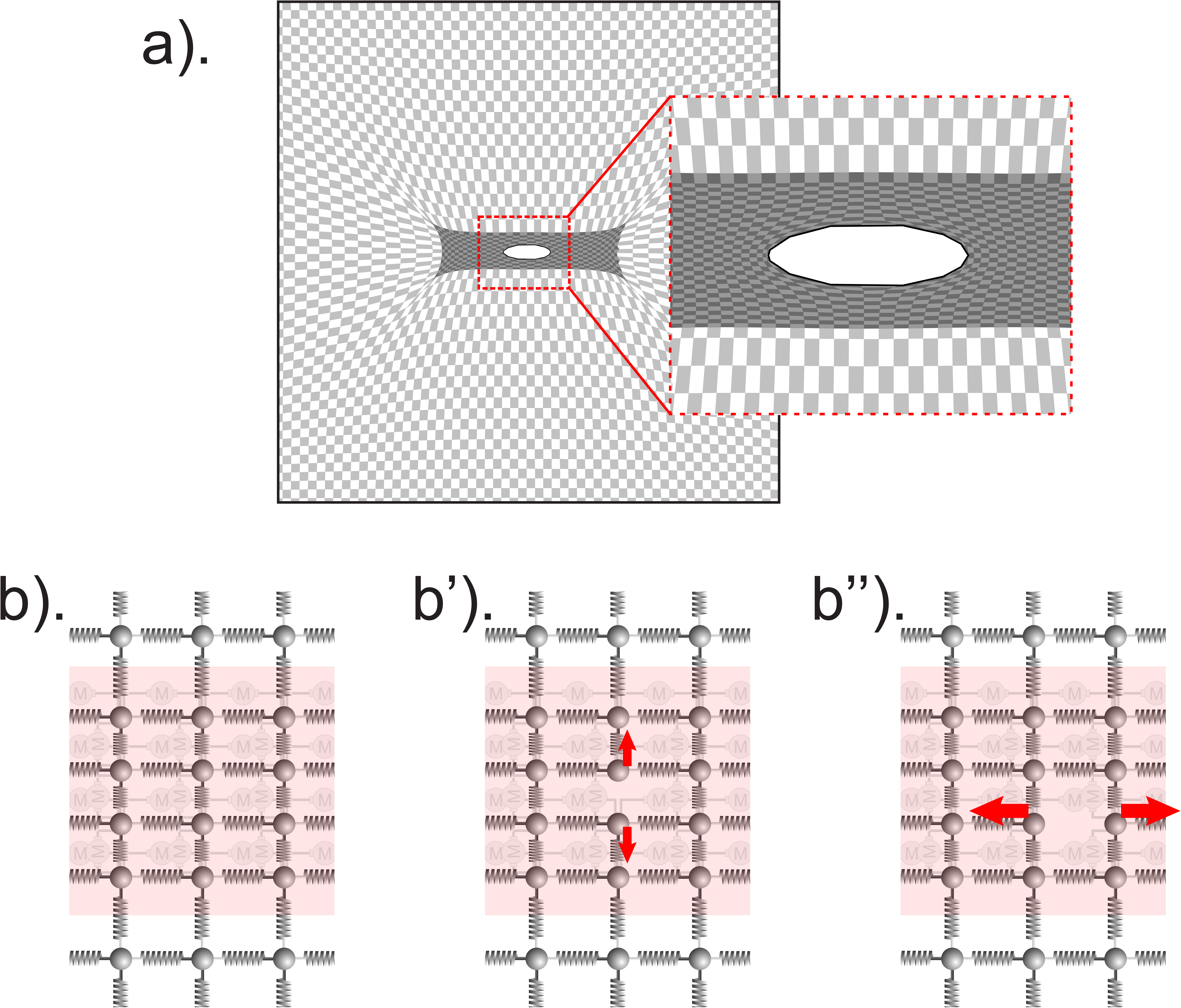
Simulation of an ablation experiments. a). Final state following a simulated laser ablation. A set of vertices in the center of the contractile domain are removed (“ablated”) after a finite deformation sets in. Parameters and the details of implementation are given in Appendix. b). Schematic intuitively explaining the outcome of the simulation in a), see main text for the details.

In order to better understand the dynamics of the minimal model in simpler terms without resorting to complex simulations, we analyze Equation (1) analytically. Firstly, it is readily seen that Equation (1) is linear and can thus be solved using Fourier transform. After a somewhat lengthy but straightforward calculation, one may obtain an exact solution of Equation (1) in the form of a Fourier expansion, see Appendix A. Although this analytical treatment is useful to re-confirm some of the conclusions obtained from the numerical integration, the final expressions appear rather complicated and not particularly telling. Thus, to further analyze the dynamics of the minimal model, we resorted to the analysis of asymptotic properties of certain limiting cases. To this end, we consider a situation where the elastic sheet is infinite in both dimensions. We choose coordinate axes to lie in the plane of the (flat) elastic sheet, and the center of the contractile domain is at the origin. Finally, we assume that the length to width ratio, *L_x_/L_y_*, of the contractile region is very large. First, let us consider the dynamics of the width of the contractile domain in the vicinity of its center. In other words, let us focus on the dynamics of a stripe-shaped semi-infinite region of small length centered on the x-axis, i.e. −*ϵ* < *x* < *ϵ*, −∞*<y <*∞, ϵ << *L_x_*. To better visualize these constructions, see schematic in Figure S1. At sufficiently short times, the dynamics of this stripe-shaped region is essentially independent of the *x*-coordinate and is thus approximately onedimensional. Hence, to describe this situation one may consider an infinite one-dimensional elastic line carrying uniform contractile stress in an interval of length 2*ϵ*. Since we assumed a linearly elastic constitutive law, the corresponding one-dimensional equation takes the form of a diffusion equation with a source term that has a form of two delta peaks, positioned symmetrically around the origin. The first peak (of magnitude −*ϕ*) is positioned at *y* = *ϵ* and the second peak of the same (but opposite) magnitude *ϕ* is centered at *y* = −*ϵ*, see schematic in Figure S1. This one-dimensional model is readily solved, showing that the displacement of each peak as a function of time will initially increase as a *t*^l/2^ power law. At long times, the two delta-peaks will begin to interact, ultimately meeting and stopping to move, provided stress magnitude *ϕ* is high enough. If *ϕ* is low, the final width of the contractile domain will remain finite. Next, let us consider the dynamics of the length of the entire contractile domain. For simplicity, let us suppose that contractile active stress *ϕ* is sufficient to shrink the shorter vertical dimension of the contractile domain to zero after finite time. After this time the horizontal dimension of the contractile domain will still be non-zero provided the aspect ratio of the domain is chosen to be sufficiently large. Following complete collapse of the width of the domain, we are essentially left in a situation where the dynamics are driven by two delta-peaks centered at the left-most and the right-most boundaries of the contractile domain. In other words, the left-most and the right-most boundaries of the domain behave like two point dorces within a two-dimensional elastic continuum (see Figure S1, left panel). It has previously been shown (see [10]), that the displacement of a region of a two-dimensional elastic continuum subjected to a point force increases logarithmically in time. In this sense, contraction along the length is much slower than along the width at short times.

The previous paragraph discussed the transient dynamics in the limit of large length to width ratio. What about the asymptotic “equilibrium” configuration? Our simulations show that the asymmetry persists after mechanical equilibrium is reached. It does not appear straightforward to give an intuitive account for this conclusion. It is also important to notice that if the contractile domain is not embedded in elastic continuum, the equilibrium configuration will not be anisotropic, as may be seen by directly solving Equation (1) analytically (to this end it suffices to assume that the solution is linear, i.e. *u_x_* = *b_x_x, u_y_* = *b_y_y*, with *b_x_,_y_* - constants, and use the boundary conditions). Thus, the conclusion that the deformation remains anisotropic at long times appears a consequence of the particular geometry in our problem. Note, however, that the main effect described in our model is a special case of a more general phenomenon when an (elastic) solid body is subjected to uniform isotropic stress: when mechanical equilibrium sets in, total stress needs not remain either isotropic or uniform. A further discussion of a similar effect in the context of a different developmental model, *C. elegans*, may be found in the following recent article [12].

To summarize, Equation (1) provides a simple model that can explain the asymmetric constriction of the mesoderm in *Drosophila melanogaster*. As was mentioned above, the same model can explain the apparent anisotropy observed in laser ablation experiments. To see this, we performed simulations where upon the onset of the constriction of the contractile domain, we removed a subset of vertices located in a round domain in the center of the contractile region (to be more specific, the ablated region is round in its deformed, not its undeformed state). Following this, the resulting “hole” opens up more in the direction perpendicular to the axis of contraction (i.e. the shorter dimension of the contractile domain). Intuitive account for this observation was given above.

## 4 Discussion

In this paper we considered a simple model that can explain the asymmetry of contraction of the mesoderm in *Drosophila melanogaster*. In our model, the asymmetry of contraction is a consequence of the asymmetry of the geometry of the contractile domain: the domain contracts along its shorter dimension *because* it is shorter along that dimension.

Let us now compare and contrast our view with a number of alternative models. One alternative possibility is that active stresses in the contractile domain are anisotropic, i.e. the cells have a preferred axis and “know to” contract along the corresponding direction. We believe that this scenario is unlikely. Firstly, there is no evidence of cell polarity in mesodermal cells of the prospective ventral furrow in *Drosophila melanogaster*. Moreover, this interpretation is at odds with the observations by Lepin and Roth, where it was shown that contractile grafts of mesodermal tissue contract along their shorter dimension regardless of their position or orientation within the embryo. If active stresses in those contractile patches were anisotropic, there has to exist a mechanism to align those stresses with the shorter axis of the domain through some self-organized process. Although not impossible, this appears a much more complicated explanation than the one we propose. Additionally, this alternative model would imply that the anisotropy as measured by laser ablation experiments would be present immediately after the onset of contraction, rather than building up gradually, as has been shown experimentally.

Another model could hold that material properties of the contractile domain are anisotropic. For instance, one can imagine that the cells of the contractile domain are easier to contract along the mediolateral axis than along the anteroposterior direction. This possibility may be refuted on the same basis as the possibility of anisotropic active stresses. Yet another possible explanation is spatial nonuniformity of active stresses. For instance, much like in the model proposed in e.g. [13, 14], one can imagine that active stresses are isotropic but form a gradient that peaks at the center of the contractile domain and decays gradually along the mediolateral direction. In this case, it may be shown that contraction will be directed along the medio-lateral axis of the embryo. However, this scenario is at odds with the results by Leptin and Roth. In these experiments, the patch of mesodermal cells descended from a small number of transplanted nuclei and it seems highly unlikely that any gradient would be preserved in this process; nevertheless, the patch consistently constricts more along the shorter axis. Additionally, we should like to stress that all the essential ingredients of our minimal model have been established experimentally. Notably, in our recent experimental study, using direct rheological measurements, we have demonstrated that embryonic tissue is highly elastic and that elasticity persists on a time-scale that is well comparable to the time-scale of gastrulation [10]. More specifically, we obtained nine minutes as a lower bound on the time-scale of elastic stress relaxation which is about the same time as that required for the mesoderm to completely contract prior to the onset of invagination.

As was detailed above, our model provides several “postdictions” about the behavior of the mesoderm. We shall like to expand on this with some predictions. Firstly, suppose that active stresses in the contractile domain decay abruptly at the boundary of the contractile domain. Suppose one measures the displacement of the different points along the mediolateral axis of the embryo. If stresses are indeed discontinuous along the mediolateral direction, one would expect the derivative of displacement measured as a function of length along the mediolateral axis to exhibit a discontinuity as well. This may readily be done with live imaging, although, we believe, such an experiment would put fairly high demands on the spatio-temporal resolution. Suppose now that one constructs a detailed quantitative model of the material properties of embryonic tissue. Such a model may be determined through rheological measurements of embryonic tissues where one subjects the tissue to controlled forces to record the resulting deformations. Our recent work [10] provides initial steps in this direction. Assuming stresses are uniform in the contractile domain and zero outside, the knowledge of the material properties of the tissue will enable to (without fitting) uniquely determine the dynamics of the mesoderm contraction. Additionally, it may be interesting to more systematically study how the geometry of the contractile domain influences the geometry of its contraction. This may be done by means of recently developed optogenetic methods allowing one to manipulate active forces involved in driving gastrulation with a high degree of control [15].

To summarize, this paper presents a simple model that can account for the asymmetry of mesoderm contraction. In this model, the asymmetry of contraction arises from the asymmetry of the domain, i.e. one side of the domain being longer than the other. This behavior relies on certain assumptions about the material properties of embryonic tissue. In particular, in order for the proposed model to describe experimental observations, the tissue must be elastic as has indeed recently been demonstrated by direct measurements.

## A Analytical treatment

In this section we solve Equation (1) using Fourier transform. We will be considering Equation (1) on a square domain with side length L. Rectangular contractile region *x*_0_ *< x < L − x*_0_*, y*_0_ *< y < L − y*_0_ is assumed to be under constant isotropic stress of magnitude *ϕ*. We Fourier-expand **u** = [*u_x_*(*x, y*)*,u_y_*(*x*, *y*)] as

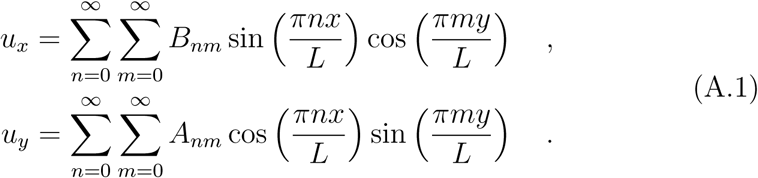

It is useful to introduce simplified notation *α ≡* E/(2(1 + *σ*)), *β* ≡ *E*/(2(1 − *σ*)). Substituting (A.1) into Equation (1) and collecting coefficients corresponding to the same modes we obtain

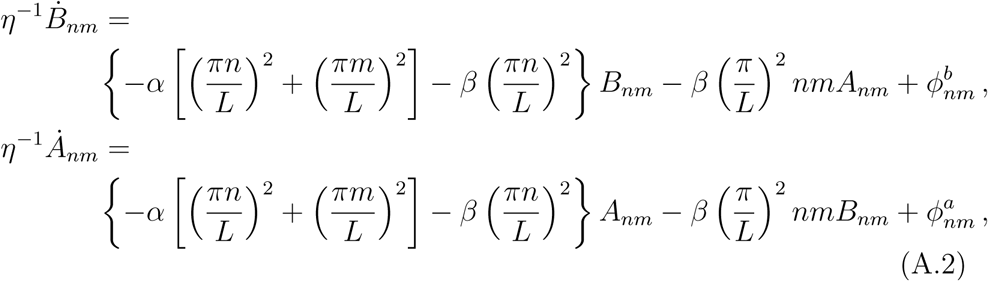

where the constant coefficients 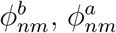 are given by

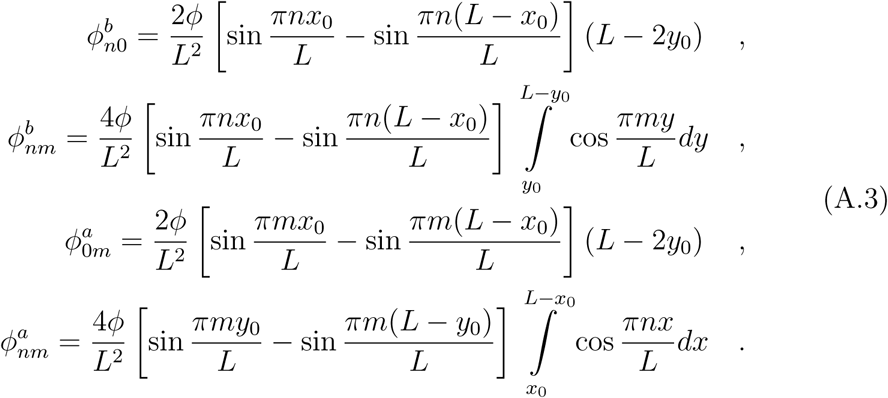

With the obvious identifications, Equations (A.2) have the form

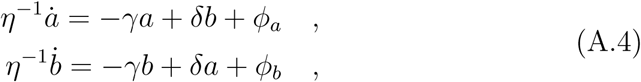

which is readily solved as

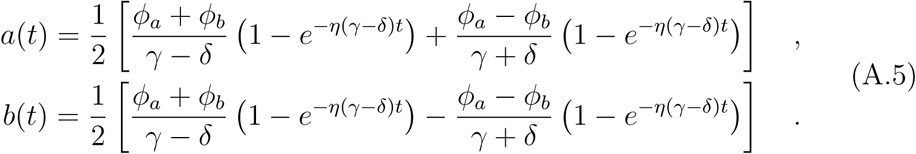

Combining Equations (A.1)-(A.5), one obtains an explicit expression for displacement **u** as a function of time and position. Since the complete expression is rather lengthy and not especially telling, we do not give it here. Instead, we present plots showing solution **u** obtained by directly evaluating the (truncated) Fourier-expansion at different time-points in Figure 2.

## B Numerical Simulations

Simulations in Figure 2b were implemented as follows. All spatial derivatives were approximated as second order central differences. The displacement of the (outermost) boundary was initially set to zero and not updated (fixed boundary conditions). Since the deformation remained localized away from the boundary throughout the course of the simulation, the particular choice of the boundary conditions is expected to not influence the result. Time integration was done by Euler-forward (explicit) scheme. The domain was discretized into a regular grid of size 800x800; time-step was *dt* = 10^−5^. In order to avoid development of singularities, we “smeared out” the forces at the boundaries of the contractile domain over a region of finite size. Specifically, force distribution had Gaussian profile exp(−*r*^2^/*ξ*^2^), with *ξ* = 0.3 (this was however fount to net be necessary as no singularities developed in the limit of much smaller *ξ*). Parameters are listed in the corresponding figure caption.

Figure 2a was generated by evaluating the Fourier-expanded expressions given in the previous section. 600 lowest Fourier modes were summed. We checked explicitly that analytical results closely matched the results of the numerical simulations.

Figures 2c,d were generated by evaluating the Fourier-expanded expressions (as was done for Figure 2a) for different choices of time and Poisson’s ratio.

Simulations in Figure 3 were done using a numerical scheme that differed from simulations in Figure 2a in order to simplify the implementation of ablation. Specifically, instead of using a finite difference scheme, we discretized the domain as an (unstructured) set of equilateral triangles whose edges are linear springs. It has been shown that this approximation reduces to the equations of linear elasticity (i.e. Equation (1)) in the limit of small strains, see e.g. [16] and [17]. Simulation parameters were chosen as follows. The simulated domain had a size of 2×2, the length of an individual spring edge was 0.027. Stiffness of an individual spring edge was set to 50. Contractile domain had a length of 0.8 and a width of 0.4, and was positioned in the center of the simulated domain. We used fixed boundary conditioned (as in Figure 2). Ablation was simulated after *t* = 0.5 by effectively removing all nodes and edges within a circle of radius 0.025 in the center of the domain. Integration was done using Euler-forward scheme with a time step of 5 ⋅ 10^−4^.

---

**Figure S1.**
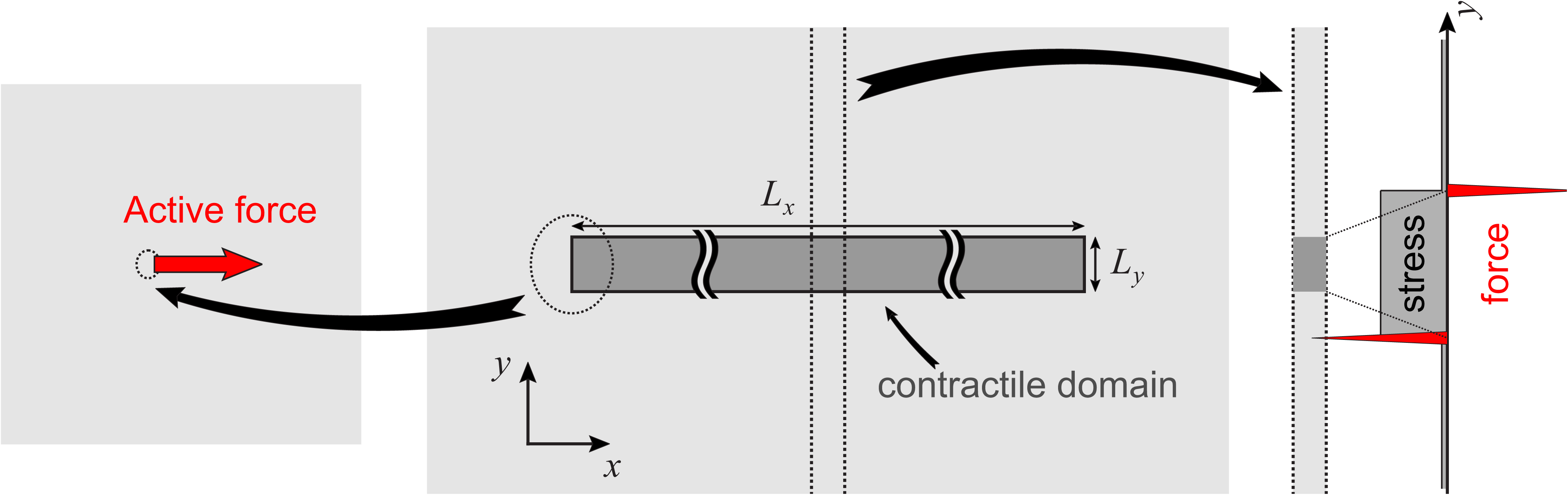
Schematic illustrating the asymptotics of the deformation in the case of very large lngth to width ratio of the contractile domain. Left pannel illustrates the dynamics of the deformaion in the vicinity of the leftmost boundary. This behavior is largely similar to that of an isolated force monopole embedded in two-dimensional elastic continuum. Right panel shows that the dnamics in the vicinity of the center of the domain is quasi-one-dimensional. See text for details.

## References

[1] Demerec M. Biology of Drosophila. Cold Spring Harbor Laboratory Press, 1994. ISBN-10: 0028438701, ISBN-13: 978-0028438702.

[2] Campos-Ortega J.A., Hartenstein V. The Embryonic Development of Drosophila melanogaster. Springer Verlag, 2013. ISBN13 9783662224915, ISBN10 3662224917.

[3] Müller H.-A. J., Wieschaus E. *armadillo, bazooka*, ans *stardust* Are Critical for Early Stages in Formation of the zonula adherens and Maintenance of the Polarized Blastoderm Epithelium in *Drosophila*. J. Cell Biol., 134, 1, 149–163.

[4] Sweeton D., Parks S., Costa M., Wieschaus E. (1991) Gastrulation in Drosophila: the formation of the ventral furrow and posterior midgut invaginations. Development. 112, 775–789.

[5] Leptin M., Grunewald B. (1990). Cell shape changes during gastrulation in Drosophila. Development. 110, 73–84.

[6] Lepin M. *twist* and *snail* as positive and negative regulators during Drosophila mesoderm development. Genes Dev. 1991 5, 1568–1576.

[7] Martin A., Kaschube M., Wieschaus E. (2009) Pulsed contractions of an actin-myosin network drive apical constriction. Nature. 457, 495–499.

[8] M. Leptin and S. Roth. (1994). Autonomy and non-autonomy in Drosophila mesoderm determination and morphogenesis. Develoment. 120, 853–859.

[9] Martin A.C., Gelbart M., Fernandez-Gonzalez R., Kaschube M., Wieschaus E. (2010). Integration of contractile forces during tissue invagination. J. Cell Biol. 188, 735.

[10] Doubrovinski K., Swan M., Polyakov O., Wieschaus E. (2017) Measurement of cortical elasticity in Drosophila melanogaster embryos using ferrofluids. PNAS in press.

[11] Landau L.D., Lifshitz E.M. Theory of Elasticity (Volume 7 of A Course of Theoretical Physics) Pergamon Press 1970.

[12] Vuong-Brender T.T.K., Amar B.A., Pontabry J., Labouesse M. The interplay of stiffness and force anisotropies drive embryo elongation. https://doi.org/10.1101/095752

[13] Mayer M., Depken M., Bois J.S., Julicher F., Grill S.W. (2010) Anisotropies in cortical tension reveal the physical basis of polarizing cortical flows. 467, 617621.

[14] Behrndt M., Salbreux G., Campinho P., Hauschild R., Oswald F., Roensch J., Grill S.W., Heisenberg C.-P. (2012). Forces driving epithelial spreading in zebrafish gastrulation. 338, 257.

[15] Guglielmi G., Barry J.D., Huber W., De Renzis S. An Optogenetic Method to Modulate Cell Contractility during Tissue Morphogenesis. Developmental Cell. 35, 5, 646660.

[16] Seung H.S., Nelson D.R. (1988). Defects in flexible membranes with crystalline order. Phys. Rev. A. 38, 1005.

[17] Liang H., Mahadevan L. (2011). Growth, geometry, and mechanics of a blooming lily. PNAS 108, 55165521.

